# MCC: Automated Mass and Charge Curation at Genome-Scale Applied to *C. tuberculostearicum*

**DOI:** 10.1101/2024.11.19.624331

**Authors:** Reihaneh Mostolizadeh, Finn Mier, Andreas Dräger

## Abstract

**Background:** For many years, antibiotics reliably protected mankind against bacterial infections, including the respiratory tract colonizer *Corynebacterium tuberculostearicum*. However, the spread of antimicrobial resistance necessitates the search for new treatment options, where the microbiota may play a crucial role. One way to investigate the complex nature of bacteria and their interactions with human hosts or microbiota is through genome-scale metabolic models (GEMs). Constructing GEMs is labor-intensive and time-consuming.

**Results:** We introduce the Python package Mass and Charge Curation (MCC), which implements a new automated algorithm to facilitate mass and charge balancing—one of the most time-consuming reconstruction steps. This package manipulates reconstructions by consolidating data from multiple resources and updating the `notes`-field with relevant changes. It also generates a visual comparison between draft and curated models, ensuring high-quality metabolic reconstructions. Using MCC, we developed a metabolic reconstruction of *C. tuberculostearicum* strain DSM 44922. The model was improved based on standardization policies, resulting in a functional, well-annotated, high-quality product. We also simulated the organism’s growth in synthetic nasal medium 3 (SNM3).

**Conclusions:** The high-quality model *i*CTUB2024RM consistently resembles growth behavior under realistic conditions in an artificial human nasal environment, enhancing understanding of *C. tuberculostearicum* and its potential impact on health and disease. The curation process of this model led to the development of the MCC package, which facilitates the mass and charge balancing of arbitrary flux balance constraints (fbc) models in SBML format.

**Availability:** MCC is freely available via PIP, from https://github.com/draeger-lab/MassChargeCuration/. The model *i*CTUB2024RM can be obtained as an SBML file wrapped in an OMEX archive from BioModels Database, accession MODEL2407310001.

**Importance:** The rise of antibiotic resistance has made it essential to explore alternative treatments for bacterial infections, particularly those caused by respiratory tract colonizers like *Corynebacterium tuberculostearicum*. Understanding the metabolic behavior of these bacteria and their interactions with the human host or microbiota is crucial. genome-scale metabolic models (GEMs) are powerful tools for investigating these interactions, but they are time-consuming to build. Our new Python package, Mass and Charge Curation (MCC), automates a crucial step in the GEM reconstruction process-mass and charge balancing-making it more efficient and reliable. By applying this tool, we developed a high-quality, functional metabolic model for *C. tuberculostearicum* (*i*CTUB2024RM), which provides deeper insights into the organism’s growth in a simulated human nasal environment. This work offers a foundation for future research into microbial communities and their role in human health.

## 1 Introduction

The diverse and healthy microbiota in the human body plays a crucial role in maintaining overall health. With the advent of advanced technologies, our ability to study these microbial communities with increasing precision has significantly improved. While much recent research has concentrated on the human gut microbiota, the nasal microbiota also plays an essential role in human health despite receiving less attention. Chronic rhinosinusitis (CRS), a condition characterized by prolonged inflammation of the nasal and paranasal sinus mucosa^1^, exemplifies the importance of understanding nasal microbiota. CRS is a prevalent condition that significantly impairs patients’ quality of life^2^. In the United States alone, CRS affected 29.2 million adults, accounting for 14.2 % of the population in 2004^3^, and it impacts approximately 5 % of the adult population in Western societies^2^. In addition, this condition resulted in annual healthcare costs exceeding 5.8 billion^4^. It has been widely recognized that microbiota plays a role in the pathophysiology of CRS. However, their exact contribution to the development and severity of the disease remains unclear^5^. Among the bacteria inhabiting the human nose, corynebacteria species are particularly prevalent^6^.

Among various corynebacteria, *Corynebacterium tuberculostearicum* was first described by Brown et al. and formally characterized by Feurer et al. in 2004. This corynebacterium is distinct from most other corynebacteria because they produce tuberculostearic acid. The cells of *C. tuberculostearicum* are non-motile, non-spore-forming, Gram-positive to Gram-variable, and non-acid-fast^7, 8^. In the sinuses, *C. tuberculostearicum* has been significantly enriched^5, 9, 10^ and is suspected to have pathological potential by mediating rhinosinusitis^9^. *Corynebacterium tuberculostearicum* commonly colonizes various skin environments, including both dry and moist regions^11^. It plays a role in skin inflammation and may contribute to chronic inflammatory diseases^11^. This bacterium is also multi-resistant to antibiotics^12^, complicating treatment options for infections it causes. Given its adaptability, understanding the environments in which *C. tuberculostearicum* thrives is essential, as it may play a role in maintaining microbial balance and contributing to the host’s immune defense mechanisms^11^. Although usually commensal, *C. tuberculostearicum* can act as an opportunistic pathogen, particularly in immunocompromised individuals^13, 14^. It has been implicated in various infections, including pneumonia, septic arthritis, and infections associated with medical devices^13, 14^. Understanding its metabolic capabilities is crucial for identifying factors that enable its transition from a harmless commensal to a pathogenic state. Moreover, metabolic reconstruction can help identify potential targets for novel antibiotics or alternative therapeutic approaches. This necessitates a focus on a GEM, which can efficiently identify and characterize the metabolic systems of nasal microbiota. A GEM model is constructed based on genome sequence annotation and physiological data, encompassing all the metabolic reactions within an organism and the genes encoding each enzyme.

A recent study using GEM on Corynebacterium species found that *Corynebacterium glutamicum* serves as an important platform for industrial biotechnology and environmental remediation^15^. GEMs play a crucial role in understanding cellular metabolism and phenotypes, designing mutant strains for desired products, and assessing the effects of genetic interventions and environmental changes on cellular metabolism^16^. High-quality GEMs provide deeper insights into the species itself, its interactions with other species, and offer more personalized treatment options for diseases caused by these species.

High-quality reconstructions of GEMs require extensive manual curation, which is often time-consuming. To streamline this process and reduce the need for manual work, we developed a Python module called MCC. This tool automatically curates mass and charge assignments for metabolites in a metabolic model. The algorithm behind the Python module is explained in detail to ensure transparency. The module gathers mass and charge information from various databases, including Biochemical, Genetical, and Genomical (BiGG)^17^, MetaNetX^18^, Kyoto Encyclopedia of Genes and Genomes (KEGG)^19^, BioCyc^20^, and Chemical Entities of Biological Interest (ChEBI)^21^, and evaluates the results. If any changes are made to the chemical formula or the number of protons in a reaction to balance the mass, these changes are documented in a `notes`-field within the model. Additionally, the module can generate a visual comparison between the draft and curated models. This allows users to see all metabolites with incomplete information or where the assignment differs from the draft model. The MCC package is freely available through a GitHub repository (see Availability, page 14).

The development of this package was driven by the goal to curate the genome-scale metabolic network reconstructions (GENRE) of *C. tuberculostearicum* strain DSM 44922. To meet the latest community standards, this reconstruction process followed best-practice recommendations by Carey et al. and adhered to the FAIR data principles (findable, accessible, interoperable, and reusable)^23^. As a result, the GEM includes a variety of features, such as fully annotated metabolites, reactions, and genes with gene-protein-reaction associations (GPRs), Systems Biology Ontology (SBO) terms^24^, Evidence and Conclusion Ontology (ECO) terms^25^, Systems Biology Markup Language (SBML)^26^ extension package for `groups`27, and KEGG^19^ pathway annotation. In addition, refinement steps were implemented to address energy generating cycle (EGC), redundancy, dead-end metabolites, and model extensions using other resources. These enhancements help fill knowledge gaps regarding growth possibilities in different environments like Lysogeny broth (LB), M9 minimal medium (M9), and SNM3^28^. The new high-quality GEM for *C. tuberculostearicum* strain DSM 44922 is available in BioModels, providing a comprehensive description of the model.

## 2 Material and methods

To achieve a high-quality GEM, it is essential to use the latest *in silico* methods and tools. Following the most current standards in systems biology, based on the reconstruction protocol from 2010 by Thiele et al., ensures the model meets these standards.

### 2.1 Reconstruction

The model reconstruction process was broken down into three main steps:: (i) creating a draft model using an open-source and user-friendly tool, (ii) manual curation for improving the model, and (iii) the analysis and quality control of the model.

#### 2.1.1 Draft model creation

The model was created using a top-down reconstruction approach with CarveMe^30^. We build on a universal model since our organism seems Gram-variable. CarveMe uses information from the universal model to *carve away* any reactions unlikely to occur in the organism’s proteome.

#### 2.1.2 Model refinement

The model created by CarveMe was based on a curated universal model using the BiGG Models Database^17^, which includes data from several high-quality, manually curated GEMs. To manipulate chemical formulae and charges of metabolites, we used the SBML Level 3^31^ fbc extension package^32^. Manual curation was performed using Constraints-Based Reconstruction and Analysis for Python (COBRApy)^33^ and the libSBML^34^ Application Programming Interface (API) for SBML^26^ in Python to improve and fill in any missing information.

##### Database annotation for reactions, metabolites, and genes

Several database annotations, including KEGG^19^, BioCyc^20^, ChEBI^21^, ModelSEED^35^, and MetaNetX^18^, were extracted from the `notes`-fields using libSBML^34^. This process standardized the data to make the model more comparable across different databases, providing a consistent way to track provenance and enabling reuse through uniform formats. This improved the reproducibility and comparability of GEMs across various databases. All annotations were added as controlled vocabulary (CV) terms with the biological qualifier type `BQB_IS`, using https://identifiers.org through the Minimal Information Required In the Annotation of Models (MIRIAM) registry^36^. This approach ensures the annotations remain accurate even if databases are modified^37^. We parsed both old and new locus tags from the GeneBank file corresponding to our specific strain for gene annotation. Depending on availability, these tags were organized into a dictionary to map gene identifiers or into a table with WP Numbers and old and new locus tags. We checked which gene identifiers (IDs) were available in our model (probably “WP …”). Using this information, we annotated the model with COBRApy. To do so, we first gathered the UniProt accession IDs from the old locus tags through https://www.uniprot.org/uploadlists/. We then queried UniProt for additional IDs. With the old locus tags, we also accessed the KEGG database via the KEGG API to retrieve references to further databases. One could use conversion methods to revert to the old tags if only new locus tags were available. Finally, all gene annotations were added to the model using libSBML^34^.

##### Extend model manually using the BioCyc database version 26.1

The BioCyc database^20^ includes metabolic pathways for *C. tuberculostearicum* strain SK141 with Reference Sequence (RefSeq)^38^ assembly accession, GCF_000175635.1 and Taxon ID `id=553206`. This database is continuously updated with new biochemical information, such as from MetaCyc^39^, using both computational and experimental approaches. Additionally, the organized assembly of organism information in National Centre for Biotechnology Information (NCBI)^40^, with RefSeq^38^ assembly accession, GCF_013408445.1, is efficiently extended. As a result, a reconstructed GEM will eventually require updates. Despite the different strains available in BioCyc^20^ and our reconstructed model, analyzing these different strains can still yield additional predictive reactions. The extension process was performed as follows:

1. Created a SmartTable with the relevant information using BioCyc^20^ and downloaded it.
2. Read the tables and organized the data into dictionaries.
3. Restored information, including reaction details, Enzyme Commission (EC) numbers, reaction direction, reaction names, substrate, and substrate formulae, and mapped their BioCyc ID to the respective values. Since BioCyc SmartTables do not contain substrate charges, we initially set each charge to 0 in this stage.
4. Parsed the protein sequences to blast them against the sequence of our strain.
5. Aligned the sequences with our strain using DIAMOND^41^.
6. Evaluated the alignment results, comparing them to those in our model based on a 95 % identity threshold, and restored the results in a dictionary.
7. Extracted BioCyc genes for the reactions and compared them to those in our model.
8. Added reactions from BioCyc^20^ for which gene evidence was found but were not present in our model.

These steps create an extendable framework applicable to any strain-specific model, making it feasible to combine comparative genomics and genome-scale metabolic modeling across multiple species with reasonable effort and expertise.

##### Mass and charge balance

We extracted all chemical formulae from the `notes`-field into dedicated attributes or MIRIAM annotations within the SBML^26^ model with fbc extension^32^, created by CarveMe^30^, and assigned each species a charge of 0. While this results in a perfectly balanced model, it does not accurately reflect biological reality. Assigning the correct charge can cause issues with imbalanced reactions, and the presence of multiple formulas for some metabolites may also lead to mass imbalances. To improve the model, each metabolite should have a unique chemical formula and the correct charge, even though this can increase the number of mass imbalance reactions. A reaction is elementally and charge-balanced once the protonation states of its reactants and products are correctly determined, which may require recovering some charge information. This balancing process can lead to an infinite loop, making it one of the most time-consuming steps during manual curation. To simplify this process and resolve charge assignment and reaction balancing, we developed an automatic Python module called MCC. This module synchronizes refining chemical formulas and associated charges, ensuring balanced reactions while saving time. The details of this approach will be explained in the next section.

##### SBO and ECO terms

To introduce additional semantic information to the model, SBO terms were assigned to all genes, metabolites, and reactions using libSBML^34^ in Python. The SBO provides controlled vocabularies commonly used in systems biology^24^. By using SBOannotator^42^, we assigned: (i) The SBO term `SBO:0000243` to all genes, representing *gene*., (ii) The SBO term `SBO:0000247` to all metabolites, representing *simple chemical*, and (iii) There are specific SBO terms were assigned based on their type, for instance, *exchange, sink, demand reaction, growth*/*biomass reaction, transport* (specifically sym- and antiport reactions), *translocation* reactions, simple reactions containing the reactant and product compartments, and specifically *efflux* and *influx* reactions.

Additionally, ECO terms^25^ (classes) were included to describe the types of evidence used, from laboratory experiments, computational methods, literature curation, or other means. These terms help to track annotation provenance, establish quality control, and provide insights into the sense of “why we believe what we think we know”^25^.

##### Groups extension

We enhanced our model by grouping pathways identified via KEGG^19^. Using the SBML^26^ `groups` package through libSBML^34^ in Python^27^, we extracted all reactions annotated with `BQB_OCCURS_IN` along with their associated pathways. For each pathway, we created a group of type `partonomy` and added the corresponding reactions, indicating that these reactions are part of a common pathway.

Additionally, we included `BQB_IS` annotations and added CV-terms of biological qualifier type `BQB_OCCURS_IN` to indicate the pathways to which reactions belong. Where necessary, we used web-requests in Python to extract missing KEGG^19^ reaction identifiers from the BiGG API and then retrieved the associated pathways from KEGG^19^ via its Representational State Transfer (REST) API. These pathways were then added to each reaction annotated with a KEGG identifier.

##### Energy generating cycles

Energy metabolism is crucial to cellular biology. Insufficiently curated GEMs can lead to thermodynamically infeasible EGCs or futile cycles without nutrient consumption. Any EGC violates the law of energy conservation, compromising the reliability of model simulations. Therefore, it is essential to accurately account for the use of external nutrients to synthesize energy metabolites in species-specific GEMs. To identify if the model contains any EGCs, we used flux balance analysis (FBA) with zero nutrient uptake while maximizing energy dissipation reactions. This included reactions for adenosine triphosphate (ATP), cytidine triphosphate (CTP), Guanosine Triphosphate (GTP), uridine triphosphate (UTP), inosine triphosphate (ITP), reduced nicotinamide adenine dinucleotide (NADH), NADPH flavine adenine mononucleotide and dinucleotide, ubiquinol-8, menaquinol-8, 2-demethylmenaquinol 8, acetylCoA, L-glutamate, and proton exchange between cytosol and periplasm, as demonstrated by Fritzemeier et al. Such cycles may occur if the GEM lacks constraints on reaction irreversibility or contains erroneous reactions or co-factors. Identified EGCs were eliminated by either constraining the reaction directionality or removing the entire reaction.

#### 2.1.3 Model analysis

The model was analyzed to identify the factors and small molecules that influence the organism’s growth rate under different environmental conditions. This was achieved by maximizing the biomass composition reaction that predicts the actual growth rate.

##### Growth on varying media

To determine the biomass components needed to sustain growth^44, 45^, FBA simulates microbial metabolism under specific environmental conditions^46^, such as LB, M9, and SNM3. LB is a nutritionally rich medium primarily used for bacterial growth^47^. M9 is a minimal salt base formulation, often supplemented with amino acids and carbon sources, and is commonly used to cultivate *Escherichia coli*^48^. SNM3 is a specialized medium designed for *in vitro* testing systems related to the human nose and supports the growth of bacteria, especially those from the Firmicutes phylum^28^. FBA can also predict nutrient utilization, product secretion, pathway usage, and missing reactions in GEM networks^49^. Furthermore, FBA computes the minimal medium requirements, i.e., the minimum number of the essential metabolic reactions needed to support specific growth rates. This minimal medium assesses growth on different carbon sources^50^. Each time, the carbon source is swapped out with another available option in the model, and growth is then recalculated.

##### Model quality

Evaluating the efficiency of GEM generation, refinement, and manual curation is challenging. To assess the quality of a GEM and its ability to generate meaningful predictions, the protocol by Thiele et al. suggests using confidence scores to refine a draft model into a high-quality one. These scores are now replaced by ECO terms^25^. While advanced platforms are available to ease manual modifications, the curation process still relies heavily on manual efforts. More time spent on manual curation typically results in a higher-quality model^51^. To evaluate the overall quality of a reconstructed GEM, the open-source platform MEMOTE is used for quality control and assurance^52^. MEMOTE assigns a score ranging from 0 % to 100 %, based on a series of tests on stoichiometric GEMs^52^. This score helps quantify the model’s quality and transparency improvements.

#### 2.1.4 Software requirements

The MCC algorithm was implemented in Python (version *>* 3.8). We also use the z3 sat-solver (https://github.com/Z3Prover/z3) to identify minimal unsat cores. All requirements, installation, and usage instructions are in the ReadMe file within the MCC project’s repository (see Availability, 14).

## 3 Results

The results here are divided into two parts; one focusing on the algorithm behind the Python module MCC for automated mass and charge curation, and the other on the reconstruction steps for model *i*CTUB2024RM strain DSM 44922.

### 3.1 The mass and charge curation algorithm

The chemical formula for the metabolites were initially originated from the `notes`-field of the draft model created by CarveMe, which pulled from BiGG Models Database^17^. However, BiGG contains some metabolites with multiple chemical formulas, missing information, or undefined side-groups (e.g., alkyl groups -R). Similar issues were found with the charges associated with these metabolites during structural reconciliation. These inaccuracies strongly affected the mass balance of reactions, leading to imbalances or inconsistencies when metabolites with undefined side-groups were present. To address these issues, we searched across databases to find the best possible match for each metabolite’s chemical formula and charge. We then paired the affected reactions in the model with database reactions if at least one participating species was involved. Since every metabolite should have an apparent molecular formula, except for the number of hydrogen atoms, which may vary, we separated the non-hydrogen balancing from the hydrogen and charge balancing. The resulting algorithm follows these six steps:

#### 3.1.1 Data Gathering

Determine and sanitize the chemical formulas and charges found in databases. This process can be easily adjusted and extended by writing and registering custom database interfaces.

#### 3.1.2 Encoding based Satisfiability modulo theories (SMT)

SMT problems involve determining whether a given logical formula can be satisfied, meaning there is some assignment of values to its variables that makes the formula true^53^. These formulas can include Boolean logic, similar to Boolean satisfiability problem (SAT) problems, which focus on finding an interpretation that satisfies a given Boolean formula^54^. They can also be more complex mathematical structures or theories. The SMT-based encoding process involves the following steps:

a. Ensure mass and charge balance for every non-pseudo reaction in the model. Charge balance means the difference in protons must match the difference in charge in the equation.
b. Assign mass and charge values to each metabolite. For metabolites where the number of protons differs from the charge, choose the most neutral form as a representative, which will be adjusted later. Remove any rest symbols for metabolites with unresolved formulas and use “≥”-constraints. Restore any unresolved rest symbols at the end of the algorithm.

#### 3.1.3 Determine balancable (SAT) core

Each reaction that appears in an unsat core is removed from the balancing process until a fully balanced subset of reactions is identified.

#### 3.1.4 Reintroduce unsatisfiable reactions

When determining the balanceable core, reactions that could not be balanced together were removed. This often means that one or more of the reactions were actually correct. Therefore, we reintroduce these reactions and check if the model can still be balanced. The success of reintroducing reactions depends partly on the order in which they are considered. To prioritize the most promising reactions, we use the following scoring scheme:

I. Problematic reactions often force a metabolite to have an unusual formula. To address this, we allowed the reactions in the model to ‘vote’ on preferred formulae. Voting involves finding all possible balanced assignments for each reaction. Each formula assignment for a metabolite was then scored based on how many balanced assignments these metabolites with this formula assignment participated. We then assumed the reaction with the lowest score in its best-scored balanced assignment as the one most likely to cause model imbalance.
II. If a problematic reaction continues appearing, we assume it is likely causing the imbalance. We, therefore, additionally weigh the previous score by considering how often the metabolite has appeared in the output.
III. Problematic reactions might overshadow unproblematic ones, incorrectly identifying harmless reactions as problematic. To prevent removing them from the model, we track which reactions appear alongside truly problematic ones. If an overshadowing is detected, the actual problematic reaction is identified, and the ‘overshadowed’ unproblematic reactions are reintroduced into the model to see if they still appear problematic.

#### 3.1.5 Optimization of assignment

Once the most extensive set of balanced reactions is identified, the model is optimized in three steps:

I. Adherence to the original models’ formulae if reactions remain balanced.
II. Use the most detailed formula available for which the model is still balanced.
III. If the formula is not constrained, use an unconstrained representation from a database. Otherwise, use a minimal amount of atoms.

#### 3.1.6 Post-processing

I. Reintroduce rest symbols for unconstrained formulas. Here, we identify metabolites as mass-unconstrained if, for every reaction they participate, another mass-unconstrained metabolite participates.
II. Choose hydrogen/charge representations adhering to the original model when possible. If not, opt for the most neutral representation, which seems like a sensible thing similar to KEGG^19^.
III. Add protons to ensure full mass balance in the reactions.

Figure 1 summarizes the steps above, and the SMT workflow is shown in figure 2. These steps led to the creation of a generic mechanism that integrates various resources—such as databases, servers, software applications, and different services—to simultaneously correct chemical formulas, assign proper charges, and balance reactions. This mechanism is packaged into a Python module called Mass and Charge Curation (MCC), which is freely available as the Python package and can be accessed from GitHub (see Availability, page 14).

**Figure 1.**
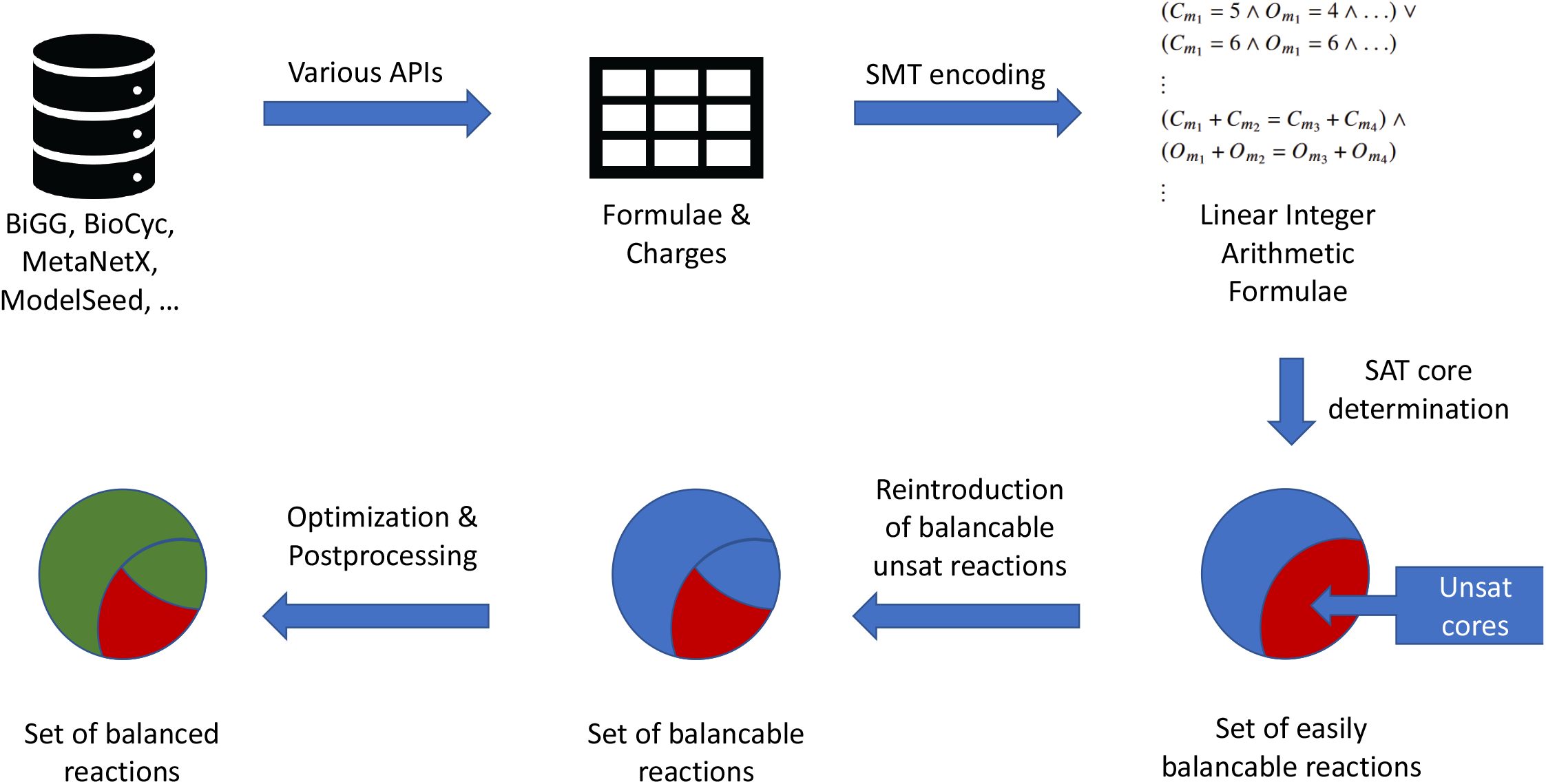
The flowchart of the algorithm used to create the Python module Mass and Charge Curation (MCC).

**Figure 2.**
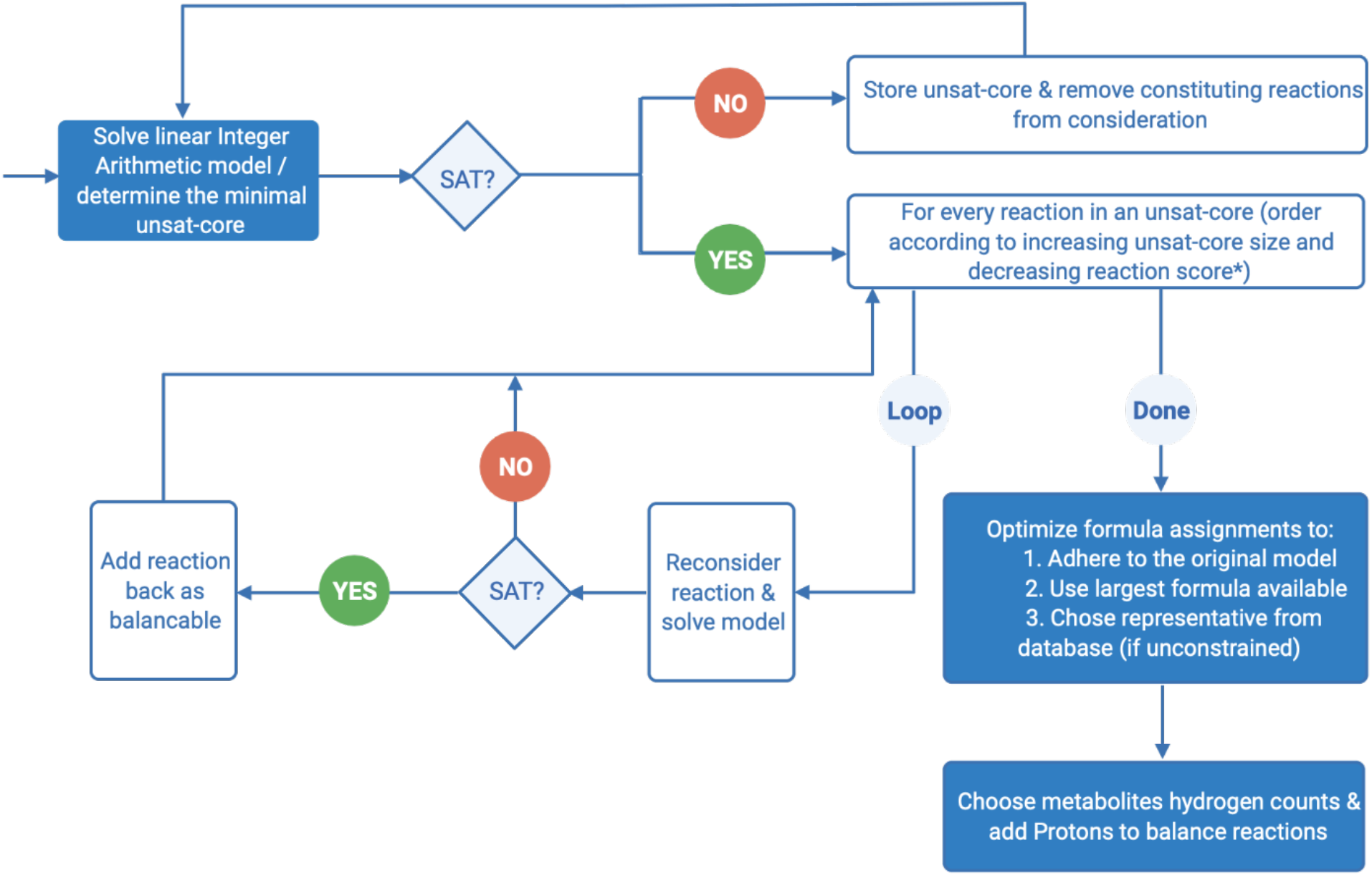
The flowchart of SMT encoding applied in the algorithm.

### 3.2 Reconstruction of *C. tuberculostearicum*

#### 3.2.1 Draft model

The first draft GEM of *C. tuberculostearicum* strain DSM 44922 was reconstructed using CarveMe^30^, including fbc package, and the available genome sequence from NCBI^40^ with RefSeq^38^ assembly accession, GCF_013408445.1. This draft served as a SBML Level 3 Version 1^55^ template. The input genome file was aligned to the universal FASTA file using DIAMOND^41^. The resulting GEM of *C. tuberculostearicum* includes 1,481 reactions, 1,019 metabolites, and is encoded by 622 genes, of which 73 are essential. The GEM identifier, *i*CTUB2024RM, follows recent community naming recommendations^22^, where *i* represents *in silico*, “CTUB” is the KEGG^19^ abbreviation for the modeled organism, “2024” marks the year of creation as iteration identifier, and “RM” indicates the lead author of this manuscript. The MEMOTE score of *i*CTUB2024RM is 38 %.

#### 3.2.2 Model curation

To curate *i*CTUB2024RM, we started by establishing GPR associations using genome annotation data, ensuring they were accurately and directly included in the model. GPR maps gene-encoding enzymes to the reactions they catalyze within Constraints-Based Reconstruction and Analysis (COBRA) models^56^. During the automated model creation with CarveMe, we used fbc to add chemical formulas for all metabolites. When multiple chemical formulas were available, we prioritized compounds with higher carbon content and without R group(s). All metabolite charges were manually assigned using the libSBML^34^ fbc package. Initially, we performed a web request from the BiGG universal model^57^ to assign these charges.

##### Metabolite, Reaction, and Gene annotation

The initial annotation of model components within the draft GEM primarily relied on information from the BiGG Models Database^17^. To enhance this, we added references from additional sources, including KEGG^19^, MetaNetX^18^, ModelSEED^35^, BioCyc^20^, ChEBI^21^, and KBase^58^. Reactions and metabolites were annotated using ModelPolisher^59^, assigning CV-Terms with the biological qualifier `BQB_IS`. These terms linked each model component to its corresponding entry in the referenced databases. For gene annotation, we utilized the GeneBank^60^ file of *C. tuberculostearicum*, extracting both old and new locus tags. We then used these tags to query KEGG^19^, UniProt^61^, and NCBI, mapping the gene identifiers to the genes in our model.

##### Model extension

Our GEM currently includes 789 metabolites involved in 954 enzymatic reactions and 63 transport reactions, all encoded by 405 genes. We compared the model to the closely related *C. tuberculostearicum* strain SK141 using data from BioCyc^20^ to refine the model. We created a SmartTable reflecting the genetic content of the target strain via BioCyc^20^ features to identify overlaps with the genomic content of our strain. Out of the 1,017 enzymatic and transporter reactions found in BioCyc^20^ for this strain, 889 were gene-encoded. We used DIAMOND for a homology search, aligning protein sequences from our strain against translated DNA sequences in the BioCyc genome database. This revealed 548 genes with a threshold of at least 95 % identity and 577 genes with 80 % identity in our model. We then mapped these gene pairs to BioCyc^20^ reactions, focusing on those with at least 95 % identity, and compared the reactions between the two strains. For each resulting gene, we checked every encoded reaction. We also reviewed all other associated genes with lower sensitivity in such reactions by involving logical expressions in their GPR. We then analyzed chemical formulas for all participating species. This resulted in a match for 192 reactions with similar metabolites, 155 of which were initially identified to be in the model through BioCyc^20^ annotation. The gene comparison revealed 205 reactions with identical genes, 140 of which were already available in the model based on BioCyc annotation. These comparisons identified 442 reactions in BioCyc^20^ with full gene evidence, 221 of which were new to our model. Therefore, the decisions are which genes possess which function, which reactions should be included, and finally, in which direction these reactions occur. As a result, we added 221 reactions and 285 metabolites to the model and modified the directionality of 355 reactions (including the added ones) based on the BioCyc^20^ data. Although these additions increased the number of orphan and dead-end metabolites, we retained them in the model to preserve its integrity as a knowledge-based model. Since MEMOTE does not account for these metabolites in its scoring, their presence does not affect the calculated score.

##### Energy generating cycles (EGCs)

Such cycles contradict the first law of thermodynamics because they produce energy-carrying metabolites, such as ATP, without nutrient consumption. To create physically reliable models, these EGCs must be eliminated. This is typically done by either constraining the directionality of reactions or removing the reaction from the network, often manually. In our model strain DSM 44922, we evaluated the production of 15 energy metabolites while preventing any nutrient uptake. When adding a reaction led to an EGC, we first tried constraining its directionality and, if necessary, removed it. However, constraining directionality can be problematic, as it may force reversible reactions to become irreversible, which is a known disadvantage^62^. To resolve all EGCs in the extended model *i*CTUB2024RM, we ultimately had to make the reaction `GLYCL` irreversible.

##### Blocked reactions

We made further improvements to identify blocked reactions in the *i*CTUB2024RM. Blocked reactions display a steady-state flux other than zero under a given medium. This can be determined using flux variability analysis (FVA) to find fluxes that carry no flux. Blocked reactions could be valuable for future refinements of the model strain DSM 44922, so we chose to preserve them in the model.

##### Mass and charge balancing

To identify the best matching chemical formulas associated with charges, correct charges associated with metabolites, and achieve mass balance in reactions whose metabolites take part in more than one simultaneous process, all reactions and metabolites’ annotations were required to be added to the SBML^26^ file of *C. tuberculostearicum* using the CV-terms. Then, our Python module MCC was applied to simultaneously correct the chemical formulas, assign appropriate charges, and balance mass in reactions by comparing data across multiple databases. This process generated a visual report (see figure 3) detailing the number of imbalanced reactions and the total running time.

**Figure 3.**
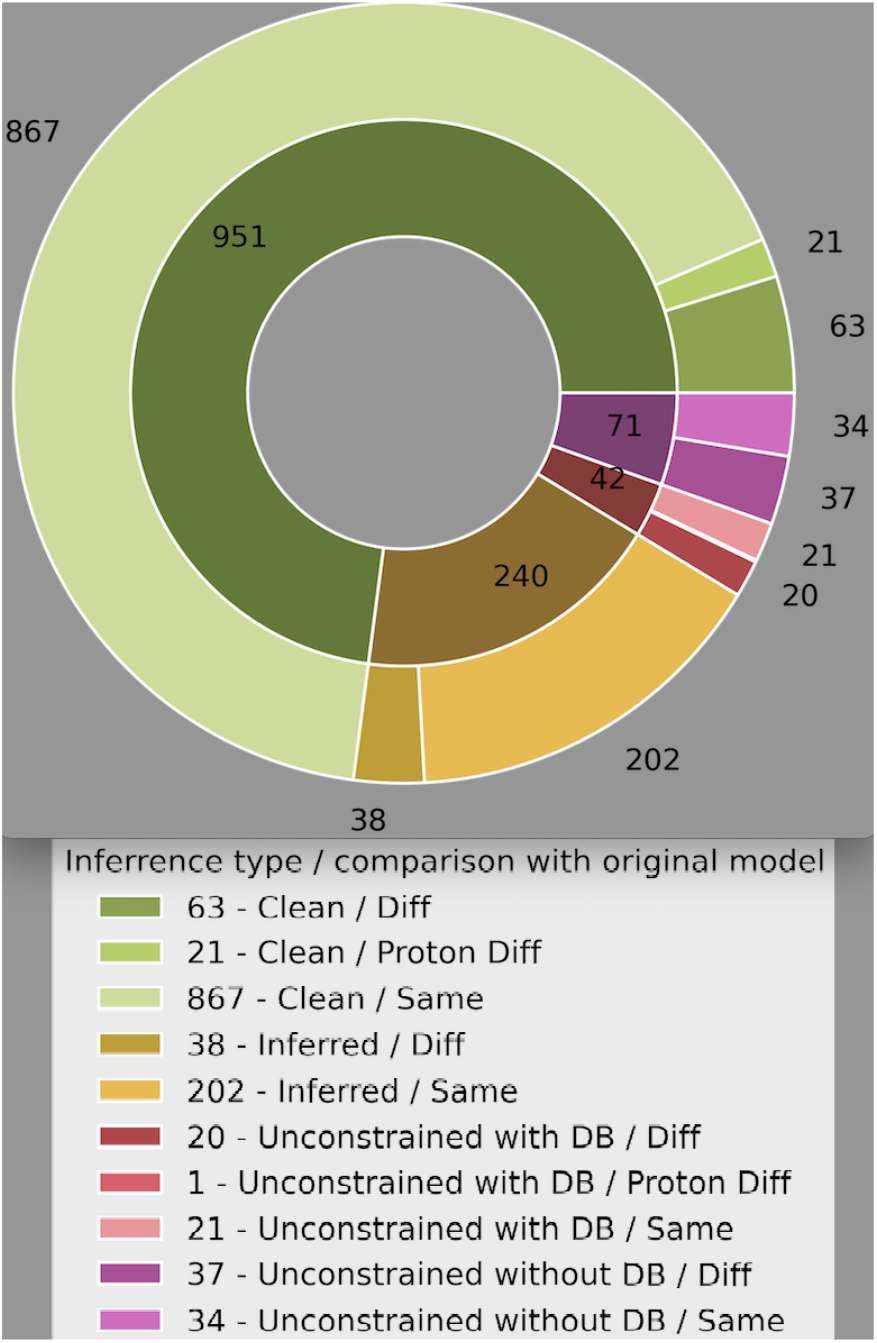
Visualization of MCC applied to the model *C. tuberculostearicum* strain DSM 44922. Metabolite formulas were compared with original formulas from 42 out of 109 initially imbalanced reactions, computed in less than 1,200 s. The pie chart categorizes the assigned metabolite formulas: Green indicates formulas from a database without undefined side-group symbols (Clean), yellow represents metabolites with formulas inferred from clean formulas and reactions (Inferred), and red/purple represents formulas still containing undefined side-group symbols (Unconstrained). The outer ring highlights deviations from initially assigned formulas, where applicable

The module’s output was summarized into reaction and metabolite reports (supplementary files 1 and 2). The relevant descriptive keywords used in these reports can be found in the tables 1 and 2. The reaction report provides detailed information about imbalanced reactions, with each table column (supplementary file 1) offering insights to help correct the model and serve as a starting point for manual curation.

**Table 1.** The relevant descriptive table of the reaction report that the module MCC produces.

| Table title | Description |
| --- | --- |
| Imbalanced Type: | This table column denotes how the reaction is problematic. This shows whether the reaction is mass or charge imbalanced or whether many protons are added to the reaction in the balancing process. |
| Reason: | For imbalanced reactions, this gives all reactions that caused this specific reaction to be excluded from our balancing efforts. The reactions listed here could not be balanced. Thus, this reaction was chosen to be taken out. In contrast, for reactions with many protons added, this denotes how many protons were added. |
| Shared Metabolites: | This column represents the IDs of metabolites that all reactions in the ‘Reason’ column share. This might likely be an indication of the problem. |
| Mass Difference: | Difference in the mass of reactants and products. |
| Charge Difference: | Difference in charge of reactants and products. |

**Table 2.** The relevant descriptive table of the metabolite report that the module MCC produces.

| Table title | Description |
| --- | --- |
| Inference Types: | ‘**Clean:**’ Formulae that came from a database and had no wildcard symbol.<br>‘**Inferred:**’ Formula for which no clean formula could be found but could be inferred from Clean formulas via reactions.<br>‘**Unconstrained:**’ Formula for which no clean formula could be found. |
| Determined formulas/charges: | This shows which formula and charge were selected. |
| Previous formulas/charges: | This shows whether the determined formula and charge are similar to one already assigned in the model. |
| Used Databases: | This indicates the determined formula was driven from which database. |
| Previous Databases: | In contrast, this shows which database supports the previous formula assigned in the model. |

Like the reaction report, we can also take a more detailed look at the formulae assigned to the metabolites through the metabolite report (supplementary file 2). Additionally, we are allowed to pass a different model to compare our assignments in this step.

As shown in the reaction and metabolite report tables (supplementary files 1 and 2), the module effectively facilitated mass and charge balancing while identifying the reactions that could not be balanced and the reasons why. It also highlighted metabolites that caused mass and charge balancing issues due to their chemical formulas or charges. The reports indicate which parts of mass and charge curation require manual attention. Consequently, starting manual curation after using the module requires significantly less effort than working directly with the draft model. In particular, formulas without ‘Used databases’ in the metabolite table are likely the most promising targets for further manual refinement. This table column helps users make additional improvements to enhance the model.

##### Redundancy in the model

First, the model was further refined by removing redundancies such as repeated reactions, metabolites, and genes. Additionally, the compartments were streamlined, retaining only three: cytosol, periplasm, and extracellular space. All identifiers in the `notes`-field were moved to the annotation section using the biological qualifier type `BQB_IS`.

##### SBO and ECO terms

SBO terms were added to the model to categorize genes, metabolites, and reactions using a controlled vocabulary. Each element was assigned a specific SBO term. ECO terms were also included during the biocuration process to describe evidence and assertion methods, categorizing them based on the functionality of gene products, such as available GPRs, spontaneous, or none.

##### KEGG pathways and grouping

CV-terms with the biological qualifier type `BQB_OCCURS_IN`, in addition to those declaring identity, were used to indicate the occurrence of a reaction within a specific pathway. These CV-terms were implemented using KEGG^19^ annotations to extract the relevant pathways.

Furthermore, the SBML Level 3^31^ `groups` extension^27^ was used to add groups for each KEGG^19^ pathway in our model, with annotations linked to the corresponding pathways via identifiers.org^36^. This allowed us to group reactions by the pathways they participate in, enhancing the model’s reproducibility and re-usability for others.

##### Growth examination

Reconstructed models may contain inconsistencies due to knowledge gaps about metabolites and reactions, which can hinder their ability to accurately predict growth under different environmental conditions. The model *C. tuberculostearicum* strain DSM 44922 reconstructed using CarveMe and refined further, successfully represented a default growth rate of 3.25 mmol/(g_DW_ · h) by setting appropriate constraints (a lower bound of −10 mmol/(g_DW_ · h) and an upper bound of 1,000 mmol/(g_DW_ · h)) for all exchange reactions.

##### Growth on SNM3

Since *C. tuberculostearicum* strain DSM 44922 has been observed in the human nose^5, 63^ it should grow on synthetic nasal medium 3 (SNM3). To simulate this growth, the exchange bounds were set to −10 and 1,000 mmol/(g_DW_ · h) except for oxygen: −20 and 1,000 mmol/(g_DW_ · h) and iron: −0.1 and 1,000 mmol/(g_DW_ · h). However, the model *i*CTUB2024RM could not produce any growth on SNM3 under these settings, so we searched for the minimal supplementation required to support growth. Adding β-nicotinamide mononucleotide (`NMN`) and cytidine-monophosphate (`CMP`) to the medium enabled growth of 1.05 mmol/(g_DW_ · h). Whether `NMN` is available in human nasal fluid^64^ remains unclear. By querying databases, we identified nicotinate `NAC` as a potential substitute for `NMN` in the human nose, which can be taken up via diffusion. We, therefore, added the corresponding reactions (as shown in table 3) to our model.

**Table 3.** Corresponding reactions of nicotinate.

| BiGG ID | Descriptive name | Reaction |
| --- | --- | --- |
| NACTpp | Nicotinic acid transport via diffusion | $\text{nac\_p} \longrightarrow \text{nac\_c}$ |
| NACTex | Nicotinic acid transport via diffusion (extracellular to periplasm) | $\text{nac\_e} \longrightarrow \text{nac\_p}$ |
| EX_nac_e | Nicotinate exchange | $\text{nac\_e} \rightleftharpoons \emptyset$ |

Breadth-first search (BFS) is an algorithm that explores nodes in a tree or graph level by level, starting from a root node. It examines all neighboring nodes at the current depth before moving to the next level, making it ideal for finding the shortest path in unweighted graphs^65^. Using this, we identified a pathway in the BiGG database that connects to one of the exchange reactions in the medium, allowing *i*CTUB2024RM to produce `CMP` in SNM3. We restricted our search to BiGG reactions with gene evidence related to `CMP`. Although we found several reactions, none could support the optimal growth objective for *C. tuberculostearicum* or cause futile cycles in the model. Therefore, the only solution was to add the exchange reaction `EX_cmp_e` to the medium.

In summary, *C. tuberculostearicum* can grow in SNM3 either by adding `NMN` and `CMP` to the medium or by incorporating the reactions listed in table 3 plus the following reaction into the model and adding `NAC` and `CMP` to the medium.

MQL8M (MQL8 Maintainance Reaction):

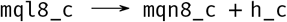

##### Growth on LB and M9

The growth of *i*CTUB2024RM was tested in LB medium. The model did not predict growth in this medium unless supplemented with `NMN`. With `NMN` added, *i*CTUB2024RM achieved a growth rate of 1.08 mmol/(g_DW_ · h). Additionally, in M9 medium, growth was supported at a rate of 0.7 mmol/(g_DW_ · h) when `NMN`, `cmp`, and β-alanine (`ala_B`) were added.

##### Minimal growth medium

We computed the minimal set of metabolic reactions required to sustain the minimum optimal growth rate for *C. tuberculostearicum*, as detailed in table 4. The growth rate achieved with this minimal medium was 0.26 mmol/(g_DW_ · h). It is important to note that this minimal medium is not unique, as it is determined through optimization. To find the minimal number of additional open exchange reactions that support a biomass objective function of 1/h, we used an optimization approach with a lower bound of −10 mmol/(g_DW_ · h) for all exchange reactions^51^. These minimal reaction sets will help examine growth rates and validate the model in future experiments.

**Table 4.**
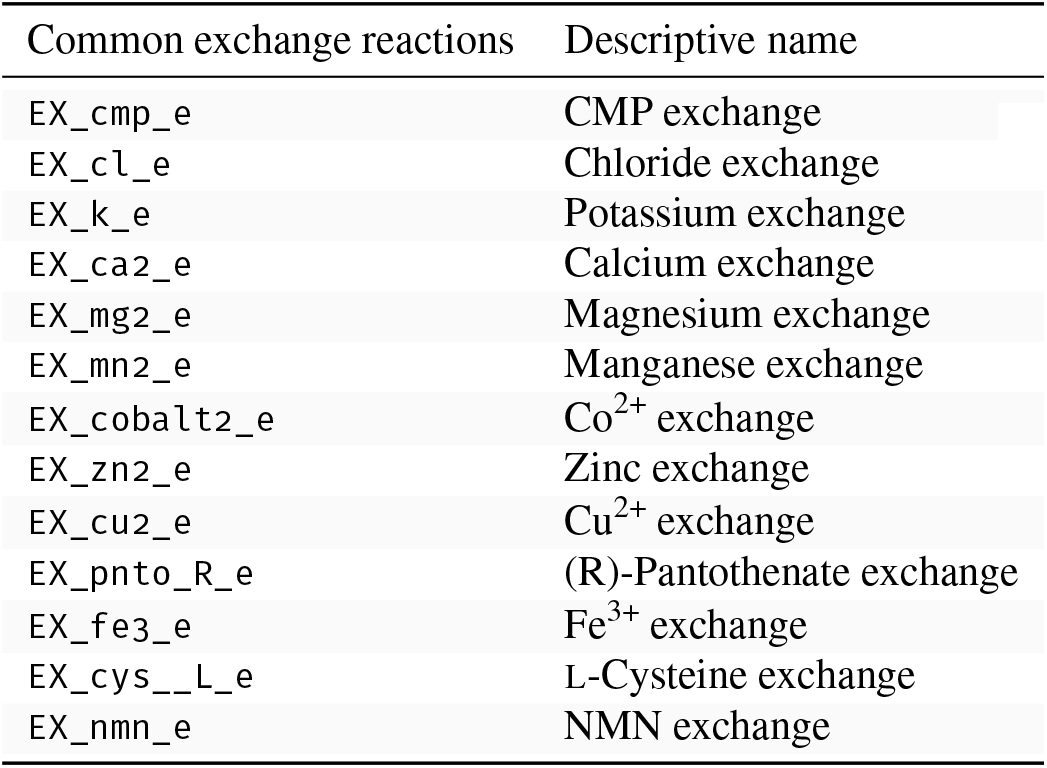
The minimal components required for the growth of *C. tuberculostearicum* with the relevant descriptive name extracted from BiGG Models Database^57^.

##### Growth on different carbon sources

To enhance the potential for experimental validation of our *in silico* predictions, we simulated the growth rate of the *C. tuberculostearicum* model using the minimal medium with different carbon sources. The minimal medium outlined in table 4, originally used L-cysteine as the carbon source. We then tested the minimal medium by replacing L-cysteine with the carbon sources listed in table 5. However, our computational analysis revealed that the *C. tuberculostearicum* strain DSM 44922 model could not grow on any carbon sources in table 5.

**Table 5.** The carbon source compounds replaced by L-Cysteine to examine the growth of *C. tuberculostearicum* with the relevant descriptive name extracted from BiGG database^57^, each causing *C. tuberculostearicum* to grow at a rate of 0 mmol/(g_DW_ · h).

| Common exchange reactions | Descriptive name |
| --- | --- |
| EX_ac_e | Acetic Acid |
| EX_cit_e | Citric Acid |
| EX_glyc3p_e | D,L-α-Glycerol Phosphate |
| EX_ala__D_e | D-Alanine |
| EX_fru_e | D-Fructose |
| EX_fum_e | Fumaric Acid |
| EX_arg__L_e | L-Arginine |
| EX_asn__L_e | L-Asparagine |
| EX_asp__L_e | L-Aspartic Acid |
| EX_glu__L_e | L-Glutamic Acid |
| EX_gln__L_e | L-Glutamine |
| EX_his__L_e | L-Histidine |
| EX_glyc_e | Glycerol |
| EX_gly_e | Glycine |
| EX_orn_e | L-Ornithine |
| EX_phe__L_e | L-Phenylalanine |
| EX_pro__L_e | L-Proline |
| EX_ser__L_e | L-Serine |
| EX_ala__L_e | L-Alanine |
| EX_glc__D_e | α-D-Glucose |
| EX_ptrc_e | Putrescine |
| EX_pyr_e | Pyruvic Acid |
| EX_succ_e | Succinic Acid |
| EX_4abut_e | γ-Amino Butyric Acid |
| EX_ile__L_e | L-Isoleucine |
| EX_mal__L_e | L-Malic Acid |
| EX_lac__L_e | L-Lactic Acid |
| EX_leu__L_e | L-Leucine |
| EX_mnl_e | D-Mannitol |
| EX_etha_e | 2-Aminoethanol |
| EX_glcur_e | D-Glucuronic Acid |
| EX_sbt__D_e | D-Sorbitol |
| EX_xyl__D_e | D-Xylose |
| EX_gly_asp__L_e | Glycyl-L-Aspartic Acid |
| EX_met__L_e | L-Methionine |
| EX_drib_e | 2-Deoxy-D-Ribose |
| EX_val__L_e | L-Valine |
| EX_malt_e | Maltose |
| EX_inost_e | L-Histidine |
| EX_rib__D_e | D-Ribose |
| EX_thr__L_e | L-Threonine |
| EX_gly_glu__L_e | Glycyl-L-Glutamic Acid |
| EX_gly_pro__L_e | Glycyl-L-Proline |
| EX_but_e | Butyric Acid |
| EX_lys__L_e | L-Lysine |
| EX_ppa_e | Propionic Acid |
| EX_man_e | D-Mannose |
| EX_tre_e | m-Tartaric Acid |

#### 3.3 Python module validation

To validate the Python module, we applied the algorithm to the GEMs of *Dolosigranulum pigrum* strain 83VPs-KB5^66^. This model was reconstructed using genomic sequence from NCBI^67^ via accession code [ASM19771v1](https://www.ncbi.nlm.nih.gov/datasets/genome/GCF_007197715.1). We used an early version of the GEM for *D. pigrum* before mass and charge balancing had been manually curated. Next, we compared the model curated using our module MCC to the published model manually curated by Renz et al. Despite manual efforts, the published model still contains 321 imbalanced reactions, including 182 pseudo-reactions, leaving 139 imbalanced reactions. As shown in figure 4, our module significantly reduced the number of imbalanced reactions, though some issues remained.

**Figure 4.**
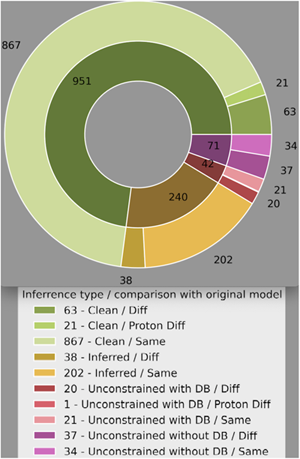
Validation of MCC applied on the model *Dolosigranulum pigrum* strain 83VPs-KB5 66 before manual curation for mass and charge balancing. The pie chart illustrates the assigned metabolite formulae in the early model compared to the manually curated version. Green represents formulae curated by MCC, showing 894 chemical formulae that match the target model exactly. In contrast, 108 are different while selected from a database without undefined side-groups symbol. Yellow indicates formulae that contain an undefined side-groups symbol inferred from clean formulas and reactions, highlighting both similarities and differences. Red/Purple denotes formulae that still contain undefined side-group symbols where a definitive formula could not be determined.

**Figure 5.**
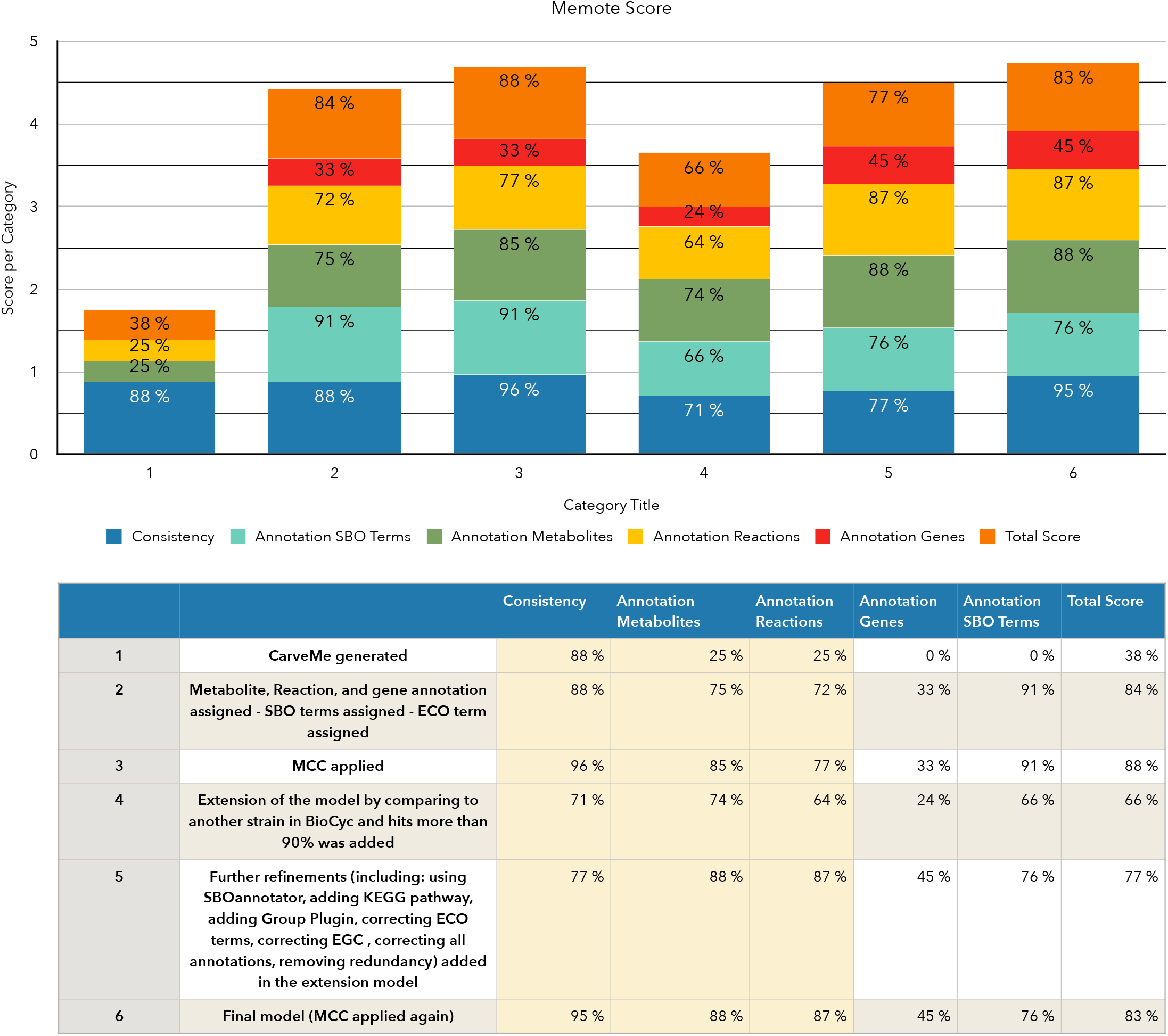
The MeMoTe scores illustrate the model’s improvement after curation and refinement. Applying MCC enhanced the model’s consistency, as MeMoTe is the result of stoichiometric consistency (SC), mass balance (MB), charge balance (CB), metabolite connectivity (MC), and unbounded flux in the default medium (UF). The score is calculated as follows: (SC + MB + CB + MC + UF) · 100/5. Although extending the model initially reduced the score due to adding reactions, metabolites, and genes, which affected mass and charge balance, further refinements improved the model’s quality. MCC helped achieve 95 % consistency and a final total score of 83 %.

**Figure 6.**
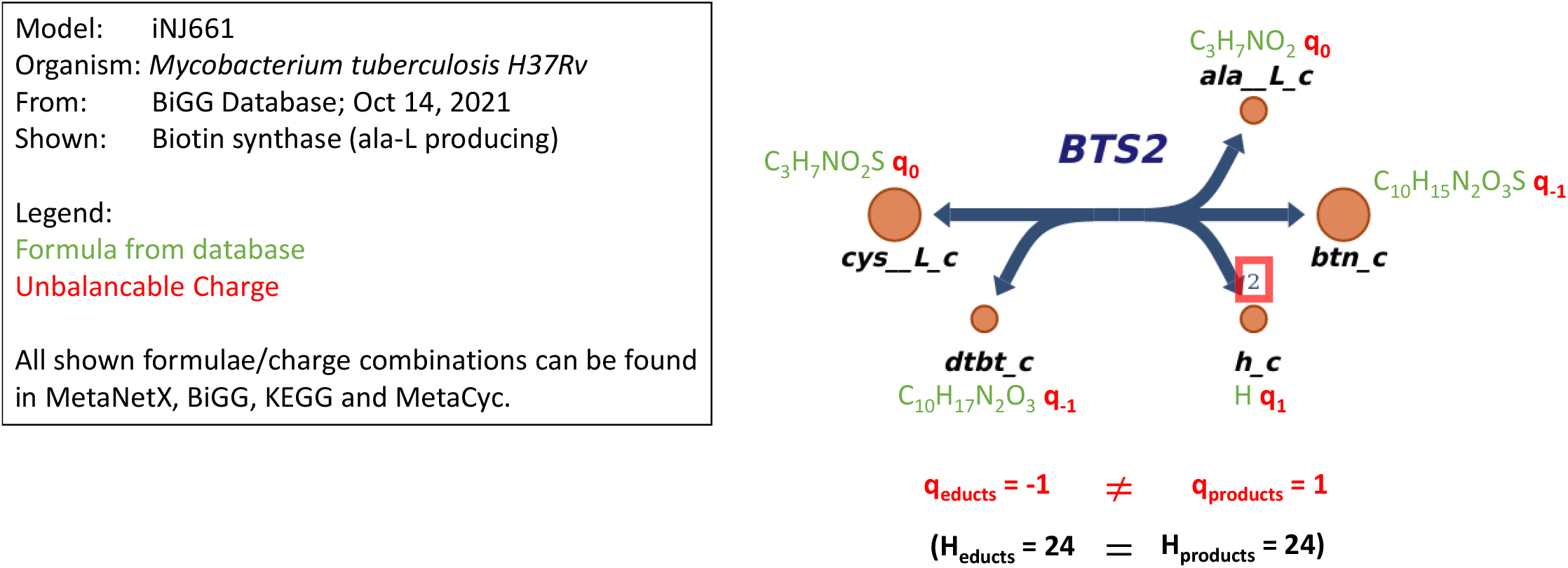
Example of an unbalanced single reaction due to incorrect charge assignments from databases.

**Figure 7.**
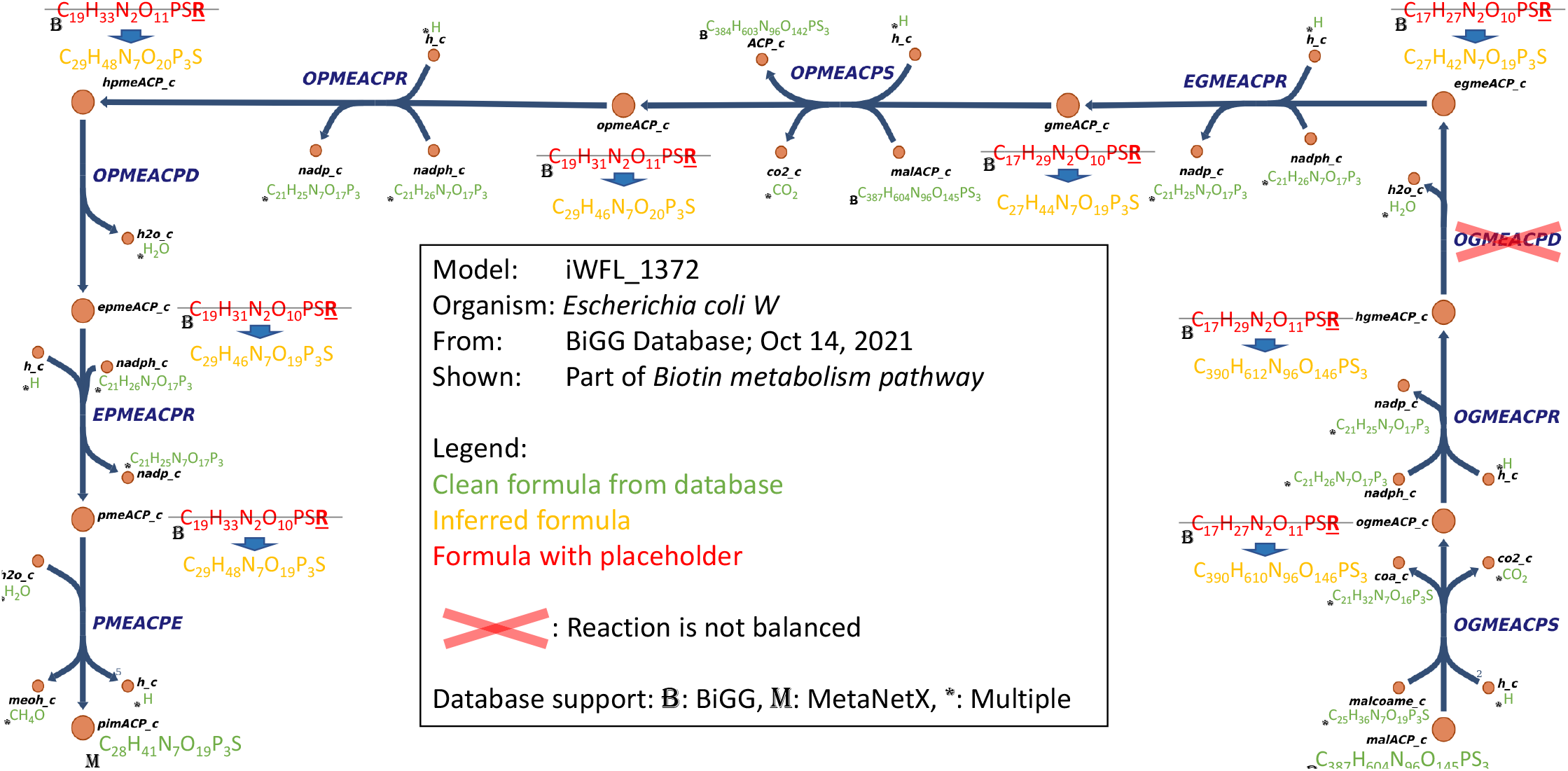
Example of an unbalanced reaction clusters and incorrect formula inference due to erroneous chemical formulae assigned by databases.

For instance, 115 metabolites were assigned differed from the target model curated manually (approximately 10 %).

These metabolite assignments were sensible in context, with most differences concentrating on metabolites that are otherwise difficult to assign manually. These include cases where metabolite annotations are ambiguous or complex, often resulting in nonsensical or overlooked assignments during manual curation. Therefore, this variation occurred because our module adheres closely to the original model and databases, leading to different decisions about balancing reactions. Additionally, the module can systematically evaluate and integrate information from various databases, while manual curation is prone to human error and limited by the curator’s ability to cross-check multiple sources simultaneously. This ensures a more comprehensive and consistent reconstruction process. These choices can vary when manually adjusting the module using the option for fixed reactions and metabolite parameters. Thus, the module not only provides more consistent results but also addresses gaps that may arise from human oversight. A comparison between models curated with MCC and those manually curated is provided in the supplementary tables (supplementary files 3 and 4). After using MCC, starting manual curation could offer significant advantages.

#### 3.4 MEMOTE comparison

An overview of MEMOTE score during the curation step is shown in 5. This indicates additional refinements enhanced the model’s quality, with MCC bringing it to 95 % consistency and a final total score of 83 %. The supplementary tables (supplementary files 1 and 2) highlight areas where manual mass and charge balancing can be further optimized. By using the following commands, the user can get the assignments that differ from the original report or/and all assignments that are not supported by any database.

metabolite_report_df[metabolite_report_df[“Similarity”] != “Same”]

metabolite_report_df[metabolite_report_df[“Used Databases”] == ““]

## 4 Discussion

Recent studies^2, 5, 9, 11, 68, 69^ highlight the significance of *Corynebacterium tuberculostearicum* in both medical and industrial fields. It plays a role in the human microbiome and can act as an opportunistic pathogen, making it important for understanding bacterial infections and developing targeted therapies. Additionally, members of the Corynebacterium genus are well-known for industrial applications, particularly in producing amino acids like glutamate and lysine^70^. While *C. tuberculostearicum* strain DSM 44922 has not been extensively studied for these purposes, understanding its metabolic mechanisms could reveal its potential and biotechnological value. To explore this, a GEM of *C. tuberculostearicum* was developed as a comprehensive tool for simulating and understanding its metabolic pathways. This model is essential for identifying the organism’s metabolic capabilities, vulnerabilities, and interactions within the microbiome, and guiding metabolic engineering in industrial processes. We initially constructed the model using CarveMe^30^, while adhering to high-quality reconstruction criteria. CarveMe^30^ typically ensures growth by adding reactions using a gap-filling approach, which means some reactions may be included that are not directly derived from genome annotations. Further refinement and manual curation were necessary to achieve accurate modeling, although one step proved particularly challenging. Manually correcting chemical formulas, charge annotations, and balancing reaction masses is complex and time-intensive. This complexity makes it essential to automate the process efficiently, especially when annotation details for metabolites and reactions are included in the model. Several software platforms have been developed to streamline this step in the reconstruction process. For instance, the SuBliMinaL toolbox^71^, written in Java™ uses third-party tools to automatically retrieve necessary data from databases like KEGG^19^ and MetaCyc^39^. It balances charge and mass by considering different stoichiometric coefficients and uses mixed-integer linear programming (MILP)^71^. Similarly, MetRxn relies on existing GEMs as references for charge calculation, using linear programming (LP) to minimize discrepancies between reactant and product charges^72^. Other approaches, like the ‘MIP’ procedure introduced by Chan et al., employ optimization techniques to resolve elemental balance inconsistencies. Despite these advancements, mass and charge balancing remains a significant bottleneck in the network reconstruction process. To overcome this, we collected mass and charge data from multiple databases (BiGG^17^, MetaNetX^18^, KEGG^19^, BioCyc^20^, ChEBI^21^) and analyzed the distribution of metabolites based on different formula-charge combinations and the support each combination received from these databases. This led to the development of MCC, a tool designed to simplify the correction of chemical formulae, charge annotations, and mass balancing in the reconstruction of GEMs. MCC is crucial because it automates a typically complex and time-consuming task, thereby improving the accuracy, reliability, and usability of GEMs for both research and industrial purposes.

This user-friendly module can be applied to any GEM. The GEM of *C. tuberculostearicum* is now available and can be used to validate growth with experimental data, study bacterial metabolism, explore pathogenic mechanisms, and guide biotechnological applications.

## 5 Conclusion

We employed systems biology to decipher the relationship between the diverse microbiota in the human nose and related diseases. Although we lack comprehensive information on the nasal microbiome and uncultured microbes *in vivo*, GEMs can grant the key steps toward understanding the principles of microbes and offer crucial insights into microbial behavior. GEMs open up a fascinating scenario of possible novel discoveries if constructed in a high-quality, manual curated manner, which is time-consuming. Recent studies have developed many automated network reconstruction tools to accelerate the reconstruction process and reduce the probability of human errors56, 74, 75, 76, 77, 78.

One of the most time-consuming steps in GEM curation is assigning the correct chemical formulae and charges to metabolites while balancing the mass in their reactions. To streamline this process, we developed a Python module called MCC that efficiently gathers relevant data from various databases and simultaneously corrects chemical formulae, charges, and mass-balancing reactions.

Our module offers a significant advantage by consolidating information from multiple sources and consistently applying it, particularly for chemical formulas. It aims to find the most accurate and consistent assignments of formulas and charges, even when dealing with conflicting resources. Additionally, the module provides visualization tools for comparing changes, allowing users to review and verify modifications rather than just accepting them. This feature also highlights areas where the module may have struggled, offering insights for further manual curation. Users who are uncertain about a formula can trace its origin and see which metabolites are affected. Unlike existing tools, which may overlook conflicting chemical formulas or refer to manual curation, our module identifies imbalanced reactions, unclear formulas, and the level of support each assignment has from various databases. An exciting feature of the module is its ability to accept a default definition by allowing users to provide a dictionary of fixed assignments for specific metabolites. For example, if a user specifies the chemical formula of water as `H2O` with a charge of 1, the module will maintain this setting during the refinement process. Unlike tools like SuBliMinaL, which balances reactions by considering different stoichiometric coefficients, our module focuses on consistency and adds protons only as needed to achieve balance. This SuBliMinaL would be more complementary than redundant with the module.

However, a limitation is that our algorithm relies solely on the balance within the model and cannot distinguish between correct and incorrect formulas if poor sources are used. With good information, though, the module enforces consistency, which enhances accuracy. Technically, the only check for accuracy is the balance, without additional *sensibility* check. Our module’s validation strategy demonstrated approximately 92 % accuracy in matching information with a well-curated model, specifically by evaluating the number of clean formulae. The validation process was straightforward to understand. We tested the MCC tool on a model before precise manual curation of mass and charge balancing. The comparison between using MCC and manual curation showed that while manual curators could achieve accurate results, they often had to stop once they reached the best possible outcome. This was likely due to the difficulty in consistently finding matching information for the remaining mass and charge balancing. Given the time-intensive nature of manual curation, our findings suggest that MCC could serve as an effective alternative, offering greater efficiency without sacrificing accuracy. For those seeking a fully balanced model, manual curation can continue after using MCC, using its suggestions for further refinement. However, achieving 100 % balancing remains challenging due to ongoing database conflicts and unresolved symbols in formulae. Despite this, MCC provides a strong foundation for the process.

Despite the advantages of using MCC, we identified metabolites involved in interconnected reactions with inconsistencies in their chemical formulas and charges. When attempting mass balancing from one side of these reactions, we encountered incompatibilities in assigning correct charges and formulas to the participating metabolites when approached from the other side. Figures 6 and 7 illustrate examples of conflicting charges and formulas, respectively.

Analyzing the nasal microbial community is a complex process with significant implications. For instance, in 1984, the bacterium *C. tuberculostearicum* was first identified in cases of diphtheria^69^. Later, it was linked to CRS^9^, showing a higher abundance in CRS patients compared to controls^5^. Recently, it has also been associated with skin cells^11^. Despite its importance in human health, the mechanisms by which *C. tuberculostearicum* influences these conditions are poorly understood. To explore the role and function of *C. tuberculostearicum* in the upper respiratory tract (URT), we first need to construct an *in silico* model of this nasal inhabitant. By using constraint-based modeling toolboxes, we can advance its application. An integrated approach in the reconstruction process could produce a high-quality model that includes elements from microbiology, ecology, and evolution.

Based on its significance in the nasal microbial community, we have constructed a high-quality GEM of *C. tuberculostearicum* strain DSM 44922. To facilitate the reconstruction process, we developed a Python module called MCC to streamline the mass and charge balancing step. We also created a script to accelerate curation for quantitative strain comparisons. Despite these advancements, comparing multiple species at the genome level^79^ still demands considerable effort and expertise. Our script can be extended to support such comparisons in other models.

We anticipate that our Python module will aid in reconstructing high-quality GEMs as more experimental data and consistent databases become available. As database conflicts are resolved, there is potential for further improving the MCC module. Future technologies will likely enhance these tools, enabling the construction of increasingly accurate models for target organisms. We hope that future studies will continue to test the hypotheses generated by the *C. tuberculostearicum* strain DSM 44922 model and provide feedback for refining the model, thus closing the loop between computational models and experimental validation and significantly reducing computational costs.

## Data availability

The genome-scale metabolic model of *C. tuberculostearicum* strain DSM 44922 is freely available from BioModels Database^80^ under accession number MODEL2407310001 as a COMBINE archive file^81^ comprising a model in SBML Level 3 Version 1^55^ format with extension package fbc version 2^32^ and annotation^82^.

While this manuscript is under review, please access the model by following these three steps:

1. Please visit https://www.ebi.ac.uk/biomodels/login/auth
2. Log in with username `reviewerForMODEL2407310001` and password `UZWSBP`
3. Access the model at this link https://www.ebi.ac.uk/biomodels/MODEL2407310001

The software for Mass and Charge Curation is freely available under the terms of the Massachusetts Institute of Technology (MIT) license from https://github.com/draeger-lab/MassChargeCuration/ or via the Python Package Index (PIP).

## Supplementary files

1. `Reaction_report_Corynebacterium_-tuberculostearicum_model.csv`
2. `Metabolite_report_Corynebacterium_-tuberculostearicum_model.csv`
3. `Reaction_report_Dolosigranulum_pigrum_-model.csv`
4. `Metabolite_report_Dolosigranulum_pigrum-_model.csv`

## Author contributions

RM and AD developed the conceptual idea. RM derived the mathematical method. FM implemented the method and algorithm under the guidance of RM. RM and FM performed the analysis and wrote the manuscript. RM and FM created the model, refined it, and analyzed it. AD updated the model and wrapped it in an Open Modeling EXchange format (OMEX) archive together with annotation and uploaded it to BioModels. AD supervised the work and critically revised the manuscript and the figures. All authors have revised, read, and accepted the manuscript in its final form.

## Acknowledgements

The authors thank Alina Renz and Lina Widerspick for providing the GEM of *Dolosigranulum pigrum* before and after their manual curation regarding mass and charge balancing to be used for the validation test of our created module in this manuscript.

## List of Abbreviations

API: Application Programming Interface
ATP: adenosine triphosphate
BFS BiGG: breadth-first search Biochemical, Genetical, and Genomical
BMBF-DZG: *Deutsche Zentren der Gesundheitsforschung*
CB: charge balance
ChEBI: Chemical Entities of Biological Interest
COBRA: Constraints-Based Reconstruction and Analysis
COBRApy: Constraints-Based Reconstruction and Analysis for Python
COMBINE: Community for Modeling Biological Networks
CTP: cytidine triphosphate
CV: controlled vocabulary
CRS: Chronic rhinosinusitis
DIAMOND: Double Index Alignment of Next-generation sequencing Data
DNA: deoxyribonucleic acid
DNA: Deoxyribonucleic Acid
DZIF: German Center for Infection Research
EC: Enzyme Commission
ECO: Evidence and Conclusion Ontology
EGC: energy generating cycle
FAIR: findable, accessible, interoperable, and reusable
FBA: flux balance analysis
fbc: flux balance constraints
FVA: flux variability analysis
GEM: genome-scale metabolic model
GENRE: genome-scale metabolic network reconstructions
GPR: gene-protein-reaction association
GTP: Guanosine Triphosphate
IBMI: Institute for Bioinformatics and Medical Informatics
ID: identifier
ITP: inosine triphosphate
KEGG: Kyoto Encyclopedia of Genes and Genomes
LB: Lysogeny broth
LP: linear programming
M9: M9 minimal medium
MB: mass balance
MC: metabolite connectivity
MCC: Mass and Charge Curation
MIRIAM: Minimal Information Required In the Annotation of Models
MILP: mixed-integer linear programming
MIT: Massachusetts Institute of Technology
NADH: reduced nicotinamide adenine dinucleotide
NCBI: National Centre for Biotechnology Information
OMEX: Open Modeling EXchange format
PIP: Python Package Index
QBiC: Quantitative Biology Center
RefSeq: Reference Sequence
REST: Representational State Transfer
SAT: Boolean satisfiability problem
SMT: Satisfiability modulo theories
SBML: Systems Biology Markup Language
SBO: Systems Biology Ontology
SC: stoichiometric consistency
SNM: synthetic nasal medium
SNM3: synthetic nasal medium 3
UF: unbounded flux in the default medium
URT: upper respiratory tract
UTP: uridine triphosphate

